# Patient-tailored design of AML cell subpopulation-selective drug combinations

**DOI:** 10.1101/2020.07.28.222034

**Authors:** Aleksandr Ianevski, Jenni Lahtela, Komal K. Javarappa, Philipp Sergeev, Bishwa R. Ghimire, Prson Gautam, Markus Vähä-Koskela, Laura Turunen, Nora Linnavirta, Heikki Kuusanmäki, Mika Kontro, Kimmo Porkka, Caroline A. Heckman, Pirkko Mattila, Krister Wennerberg, Anil K. Giri, Tero Aittokallio

## Abstract

The extensive primary and secondary drug resistance in acute myeloid leukemia (AML) requires rational approaches to design personalized combinatorial treatments that exploit patient-specific therapeutic vulnerabilities to optimally target disease-driving AML cell subpopulations. However, the large number of AML-relevant drug combinations makes the testing impossible in scarce primary patient cells. This combinatorial problem is further exacerbated by the translational challenge of how to design such personalized and selective drug combinations that do not only show synergistic effect in overall AML cell killing but also result in minimal toxic side effects on non-malignant cells. To solve these challenges, we implemented a systematic computational-experimental approach for identifying potential drug combinations that have a desired synergy-efficacy-toxicity balance. Our mechanism-agnostic approach combines single-cell RNA-sequencing (scRNA-seq) with *ex vivo* single-agent viability testing in primary patient cells. The data integration and predictive modelling are carried out at a single-cell resolution by means of a machine learning model that makes use of compound-target interaction networks to narrow down the massive search space of potentially effective drug combinations. When applied to two diagnostic and two refractory AML patient cases, each having a different genetic background, our integrated approach predicted a number of patient-specific combinations that were shown to result not only in synergistic cancer cell inhibition but were also capable of targeting specific AML cell subpopulations that emerge in differing stages of disease pathogenesis or treatment regimens. Overall, 53% of the 59 predicted combinations were experimentally confirmed to show synergy, and 83% were non-antagonistic, as validated with viability assays, which is a significant improvement over the success rate of randomly guessing a synergistic drug combination (5%). Importantly, 67% of the predicted combinations showed low toxicity to non-malignant cells, as validated with flow-based population assays, suggesting their selective killing of AML cell populations. Our data-driven approach provides an unbiased means for systematic prioritization of patient-specific drug combinations that selectively inhibit AML cells and avoid co-inhibition of non-malignant cells, thereby increasing their likelihood for clinical translation. The approach uses only a limited number of patient primary cells, and it is widely applicable to hematological cancers that are accessible for scRNA-seq profiling and *ex vivo* compound testing.

## Introduction

Acute myeloid leukemia (AML) is a heterogeneous disease, characterized by a broad spectrum of molecular alterations that influence the patient’s clinical outcomes.^1-3^ Despite the recent increase in molecularly-targeted treatment options, primary and acquired drug resistance pose a substantial challenge for most AML patients^4^. Monotherapy resistance has remained a major clinical challenge even in patients with a confirmed disease target^2,5^. For example, nearly 60% of relapsed/refractory AML patients with FLT3 mutation show resistance to gilteritinib therapy^6^. This is partly due to the activation of RAS/MAPK pathway genes through acquisition of new mutations^7^. Multi-targeted therapies provide an opportunity for synergistic inhibition of multiple resistance mechanisms, including patient-specific cancer rescue pathways and phenotypic redundancy across heterogeneous cancer subclones^8-13^. However, the emergence of resistance is a dynamic process influenced by both genetic and molecular factors, together with selective pressure from the administered therapeutics; this can be seen for example in the emergence of *NRAS*-mutated subclones in the acquired resistance to FLT3 inhibition,^7^ or intrinsic molecular and metabolic properties of monocytic subclones of patient cells resistant to venetoclax therapy^14^.

Identification of patient-specific drug combinations that target specific cell subpopulations or subclones poses a combinatorial challenge. Even though genomic analyses have led to improvements in understanding of the molecular landscape of AML, patient-specific therapeutic responses are often associated with unique clusters of non-recurrent and co-occurring mutations, making even the large-scale genomic profiling resources under-powered for identifying patient-customized combinatorial mechanisms. Furthermore, many of the most frequent AML mutations generate broad changes in the epigenome, RNA splicing and translation^15,16^, suggesting that precision medicine approaches based solely on mutation signature may fail to predict clinically useful combinations^2^. An additional clinical challenge comes from intolerable, drug-induced toxicities, especially in older AML patients^11,17,18^. The identification of both effective and safe combinations requires capturing the molecular heterogeneity of the disease progression as well as differential responses of combinations between cell subpopulations at various stages of pathogenesis, using assays that are practical for translational applications in scarce primary patient cells.

Here, we implemented a functional precision medicine approach to prioritize AML patient-specific and cell subpopulation-targeting drug combinations that uses only limited numbers of patient cells. Our mechanism-agnostic approach exploits the power of scRNA-seq technology to identify various cell subpopulations in the complex patient samples at baseline and in various disease stages. Using an efficient machine learning approach, together with compound-target interaction networks, we combined the single-cell transcriptomic profiles of the patient cells with their *ex vivo* single agent treatment viability responses to predict synergistic combinatorial inhibition of AML cells with a low likelihood for toxic effects. Using this approach, we identified both common and patient-specific synergies for 4 treatment-naïve or treatment-refractory AML patient samples, each presenting with different molecular backgrounds and synergy mechanisms. Subsequent flow cytometry experiments in the same patient cells confirmed that many of the predicted combinations led to minimal inhibition of non-malignant cells, hence reducing the likelihood of broadly toxic combination effects. To our knowledge, this is the first translational approach for systematic tailoring of personalized drug combinations that takes into account both the molecular heterogeneity of AML and toxic effects of combinations.

## Results

### Prediction of subpopulation-targeting drug combinations using limited patient cells

We applied our computational-experimental approach (**Fig. 1**) to 4 bone marrow aspirates from 3 acute myeloid leukemia (AML) patients (**Suppl. Table 1**). Similar to our previous work^19^, the computational search approach makes use of both on and off-targets of the compounds to narrow down the combinatorial search space of more than 100,000 pairwise combinations, far beyond what could be tested systematically in limited patient cells. To prioritize the top patient-specific combinations, we first compressed the genome-wide scRNA-seq transcriptomic profiles of each cell in individual patient samples into compound-target expression enrichment scores using gene set variation analysis (GSVA)^20^. The GSVA results in compound-specific enrichment scores among the cell types and protein targets of the 456 compounds tested *ex vivo* with whole-well viability assay (see Materials and Methods; **Suppl. Fig. 1)**. In the prediction step (**Fig. 1a)**, we trained a XGBoost machine learning model with conformal prediction^21^ to predict the most synergistic drug combinations for each patient sample with confidence. The model makes use of the target expression levels of the compounds both in malignant and healthy cells. This enables us to control not only the predicted overall synergistic effects, but also the predicted differential co-inhibition effects between AML and non-malignant cells.

**Figure 1.**
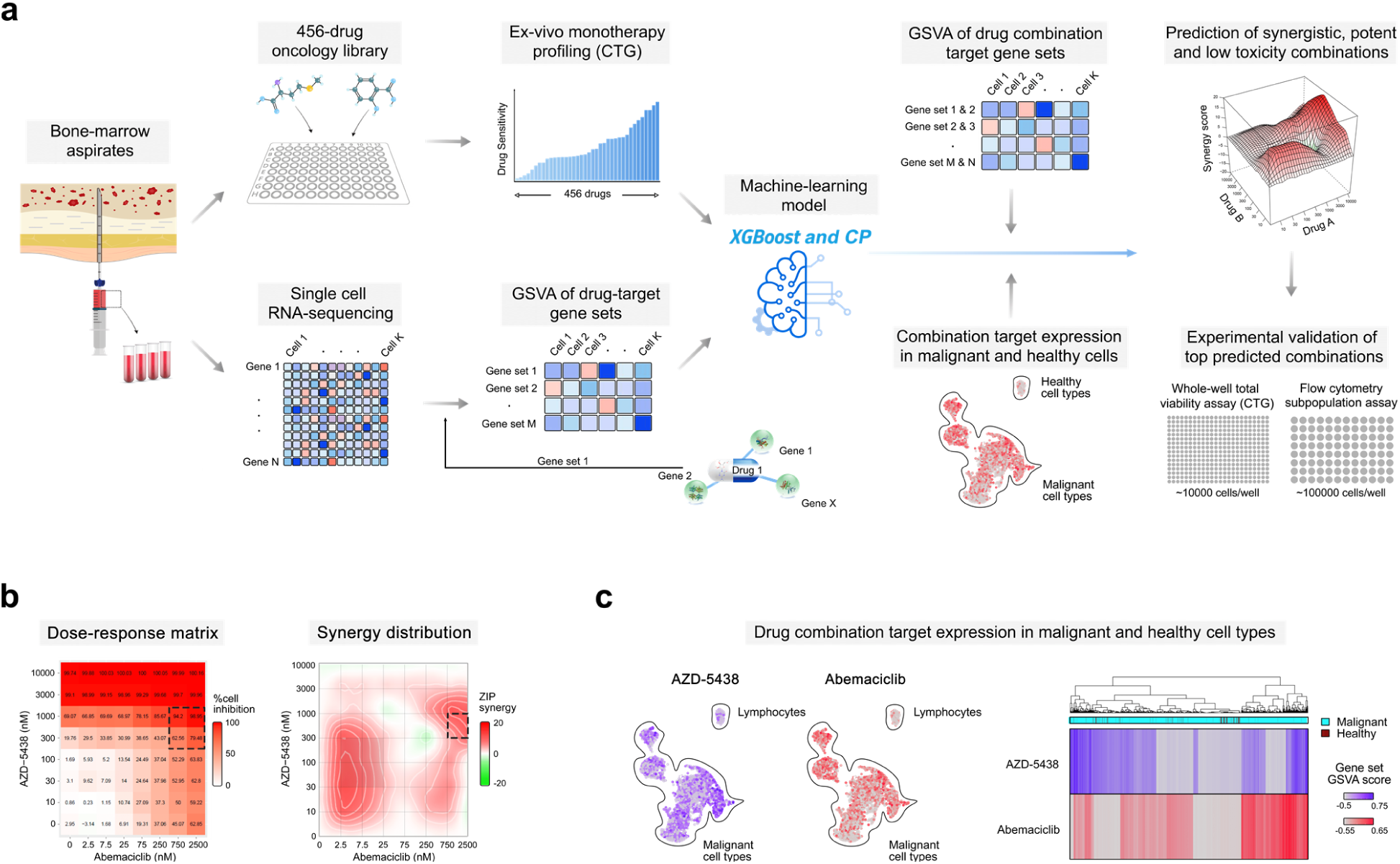
Schematic representation of the drug combination prediction approach. (**a**) Prediction of combinations with high synergy and potency in malignant cells, and low toxicity in non-malignant cells, based on high-throughput *ex vivo* single agent profiling of viability responses of individual patient sample to 456 compounds, combined with the 3’ end whole-transcriptome scRNA-sequencing of enriched mononuclear cells, compressed to compound-target enrichment score matrix (GSVA based on the single agent targets). These input data were used for training of integrated machine learning model (XGBoost combined with Conformal Prediction) for the prediction of compound-induced overall cell viability inhibition at the concentration nearest to the relative half-maximal inhibitory concentration (IC_50_) of each single agent and each patient sample separately (**Suppl. Fig. 2**). (**b**) An example of a predicted drug combination (AZD-5438 and abemaciclib) for AML2 patient sample. Combination dose-response matrix and the corresponding synergy distribution confirmed the predicted synergistic effect in the region around IC_50_ of the two compounds (dashed rectangle). (**c**) UMAP projection of the single-cell RNA-seq profiles informs about the compound-target enrichment scores across cell types of the patient sample before the *ex vivo* treatments. In this example, GSVA revealed complementary low-overlapping expression scores for the targets of the two compounds, explaining their synergistic co-inhibition effect in various populations of malignant cells. Lymphocytes were considered as non-malignant “healthy” cells in the prediction model.

In the experimental step (**Fig. 1b**), the most promising predicted combinations for each patient sample were tested in 8×8 dose-response combination matrices using *ex vivo* viability assays to confirm whether the predicted combinations truly act synergistically, i.e., jointly inhibit patient cells more than what is expected from their single agent effects, as quantified using the ZIP synergy score^22^. Importantly, after confirming the overall synergistic effects in the whole-well viability assays, we further tested a subset of validated synergistic combinations using high-throughput flow cytometry assays in the same patient cells to differentiate between combinatorial responses observed in the malignant and nonmalignant primary cell subpopulations. This experimental step prioritizes those combinations that selectively target AML cells and avoids toxic co-inhibition of lymphocytes that were considered as non-malignant cells in the prediction model^23^. The number of combinations selected for both whole-well and flow cytometry assays was based on the number of cells available in each patient sample (**Suppl. Table 1**). The combination target expression patterns from the whole transcriptome scRNA profiles provides hypotheses for molecular determinants of drug combination efficacy and toxicity in each individual patient sample (**Fig. 1c**).

### Patient-specific combinations show high overall synergy and wide inter-patient variability

We first investigated a subset of the combinations that were shared among the top-5% of predicted combinations in at least 2 patient samples. The predicted combinations resulted in co-inhibition of various targets and biological pathways (**Fig. 2a**). Two such common combinations that showed high synergy (ZIP > 5) in the whole-well viability assay across all the samples included camptothecin (topoisomerase I inhibitor) combined with etoposide (topoisomerase II inhibitor), and venetoclax (Bcl-2 inhibitor) combined with vistusertib (mTOR inhibitor). While simultaneous blocking of both topoisomerase I and II may lead to overlapping toxicities^24^, it has been shown that co-targeting of both Bcl-2 and mTOR pathway triggers synergistic apoptosis and enhances drug-induced cytotoxicity by suppressing MCL-1 in leukemic cells^25^. The predictive approach also identified the ruboxistaurin-ipatasertib combination as synergistic in all but one sample (in the refractory AML3 sample the combination was additive with ZIP = 2.3). This combination is likely to target multiple monotherapy-resistant AML subclones, as co-inhibition of PKC and AKT promotes blast cell death by downregulation of antiapoptotic Bcl-2 family proteins in leukemic cells^26,27^.

**Figure 2.**
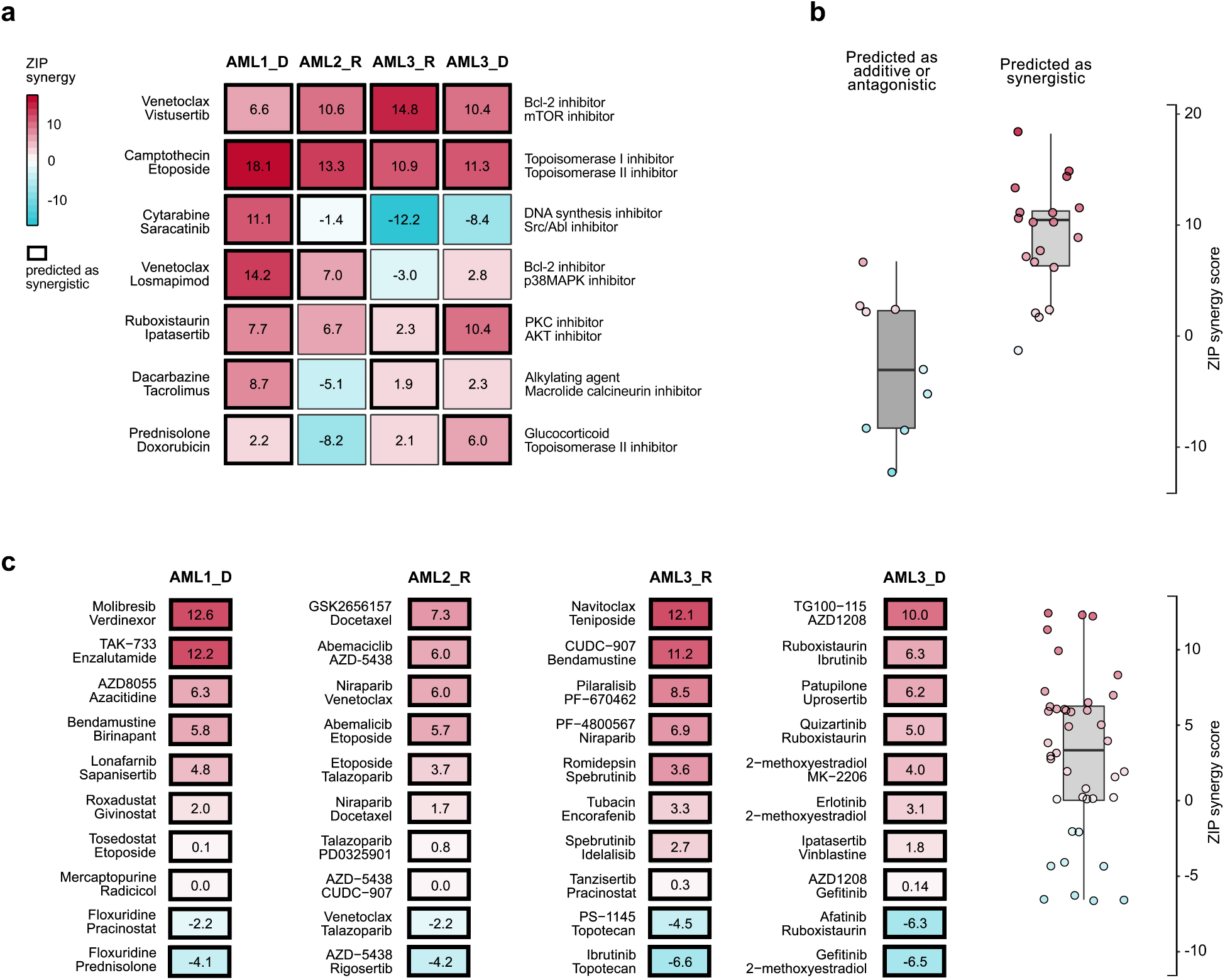
Experimental validation of the combination predictions using combinatorial CTG viability assay. (**a**) The top-7 shared combinations predicted to have synergy and AML cell-selectivity in at least two samples. The numbers correspond to the ZIP synergy score, calculated for the dose region around IC_50_ values of each drug in combination, separately for each patient sample and combination. (**b**) Comparison of the synergy score distributions of the combinations predicted to be either synergistic (ZIP>5) or antagonistic (ZIP<-5) or additive (−5<ZIP<5) in the patient samples (P<0.01, Wilcoxon rank-sum test). (**c**) The top-10 patient-specific combinations predicted uniquely for each patient sample (left), and their overall synergy score distribution in the whole-well assay (right). Overall, 53% of the 59 predicted synergistic combinations were experimentally confirmed to show synergy, and 83% were non-antagonistic (ZIP>-5).

In general, across 28 common combinations (7 combinations tested in each of the 4 samples), those combinations that were predicted to have synergy and AML-selectivity showed significantly higher synergy in the combinatorial viability assay, compared with those that were predicted to be only additive or antagonistic (P<0.01, Wilcoxon rank-sum test; **Fig. 2b**). This demonstrates the importance of patient-specificity of the predictions, even for those combinations resulting in shared synergy among multiple patient cases. For instance, we identified a strong overlapping synergy between venetoclax and the p38 MAPK inhibitor losmapimod in the two samples where the combination was predicted to be synergistic and AML-selective (AML1 and AML2), while exhibiting an additive effect in the two other patient samples. It has been shown that co-inhibition of Bcl-2 and p38 MAPK leads to synergistic decrease of phosphorylated Bcl-2, since inhibition of p38 MAPK activity alone cannot stop phosphorylation of Bcl-2^28^, causing resistance to venetoclax in leukemic cells^29^. Interestingly, AML patients with blast cells lacking Bcl-2 phosphorylation survive longer than those patients with blast cells with phosphorylated Bcl-2^26^.

As expected, the measured synergies of the patient-specific unique combination predictions were higher than those of shared combinations predicted as antagonistic or additive (P=0.03; Wilcoxon rank-sum test; **Fig. 2c**). However, the shared combinations that were predicted to act synergistically across multiple samples showed higher synergies than the patient-specific unique combinations (P=0.0002), but this difference was mainly due to the two broadly synergistic combinations, venetoclax-vistusertib and camptothecin-etoposide (**Fig. 2a**). Overall, 40% (16/40) of the predicted patient-specific combinations were experimentally confirmed to show synergy in the whole-well viability assays, when using the synergy cut-off of ZIP>5. In the 28 shared combinations, the true positive rate of the experimental validations was much higher, 79% (15/19). Among the 68 the tested combinations, there was only single synergistic combination that was not predicted by the model, resulting in <2% false negative rate. Such high precision was achieved even though the patient-specific combinations revealed a wide spectrum of co-inhibitors of multiple biological pathways active in the AML patient cells.

### Flow cytometry assay confirms cell subpopulation-specific combinatorial inhibition effects

Even though patient-specific combination designs arguably exclude broadly-toxic combinations, the whole-well viability assay cannot effectively discriminate between AML cell-killing and potential toxic effects of combinations. We therefore next investigated the degree of which the predicted combinations that showed high overall cell inhibition synergy led to co-inhibition of specific cell populations using combinatorial flow cytometry assays. To quantify the AML-selective effects, the relative co-inhibition of lymphocytes (specifically T and NK cells) was compared against the other cell populations in each of the AML patient samples separately (marked as AML cells in **Fig. 3b**). The average co-inhibition of the non-malignant cell subpopulations across the combinations predicted for the three patient cases was 40 ± 22%, significantly lower compared to the combinatorial inhibition of the AML cells (average 60 ± 22%; P=3.9×10^−6^, Wilcoxon signed-rank test). When using a cut-off of 50% for the relative inhibition of T and NK-cells, 67% (12/18) of the predicted combinations showed low toxicity, indicating the patient-specific combinations resulted in relatively selective killing of AML cells.

Notably, there were marked differences in the AML cell-selective co-inhibition effects both between the predicted combinations and patient samples. In particular, combinations involving venetoclax had relatively high co-inhibition effects on T-cells, where their potential toxic effects depended on the particular patient sample and the combination partner (**Fig. 3b**). This suggests that the immune composition of the bone marrow of these AML patients may become reshaped during the venetoclax treatment, depending on the specific combination^30^. We identified also unique combinations in single patients, such as the combination between saracatinib (SRC, ABL inhibitor) and cytarabine (DNA synthesis inhibitor), which was confirmed to be highly synergistic, low-toxic and selective in AML1 patient only. Many of these subpopulation-specific co-inhibition effects cannot be captured by the whole-well assay, and therefore required additional validation with flow cytometry assay. However, the whole-well viability assay was informative for confirming the overall co-inhibition synergy, which remained high across all the patient-specific combinations (average ZIP = 9.50 ± 3.58%; **Fig. 3a**).

**Figure 3.**
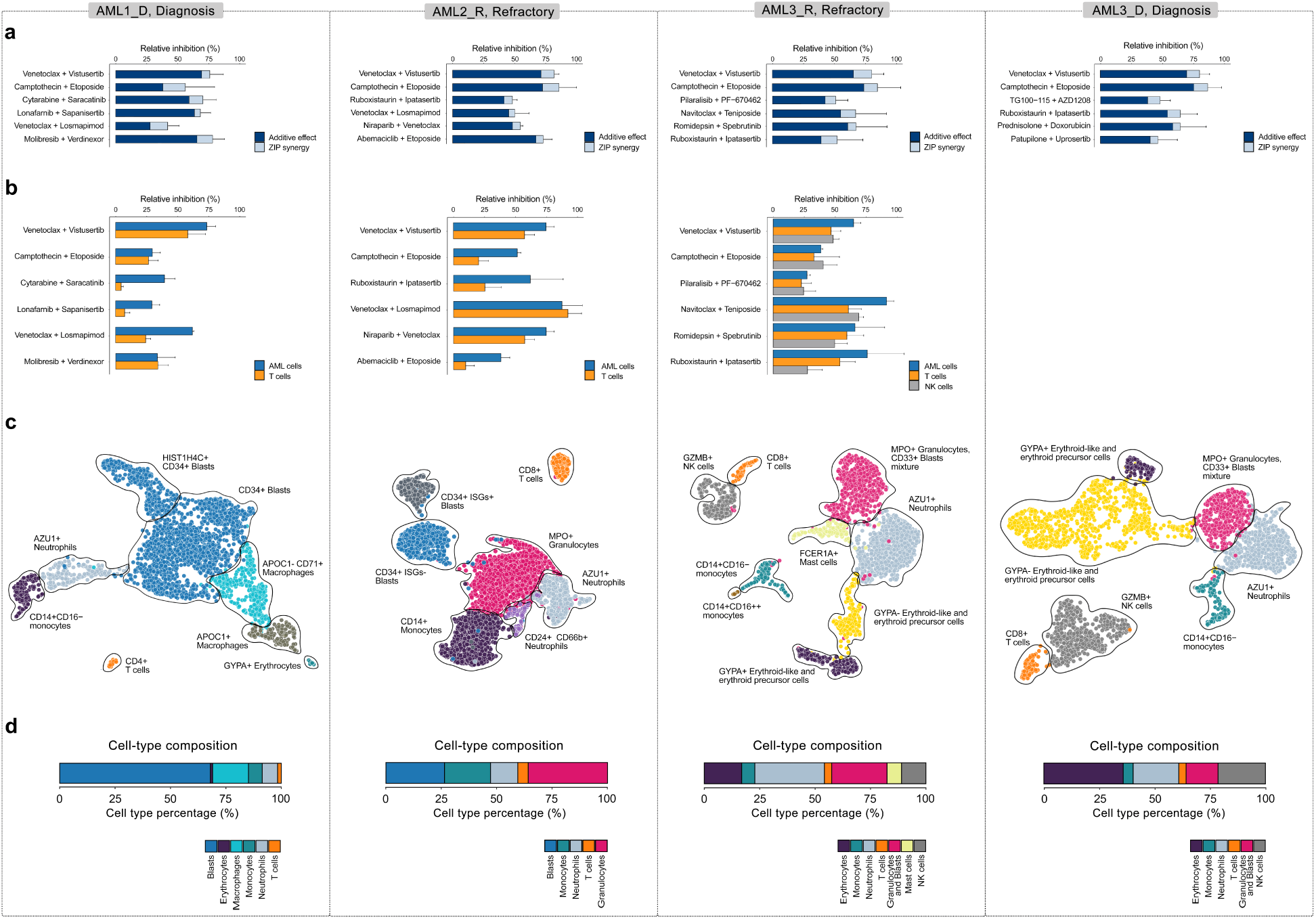
Co-inhibition effects of the predicted combinations selected for flow cytometry experiments in each patient sample (note: no cells available from AML3 patient diagnostic sample for flow experiments. **(a)** The combined dark and light blue bars together indicate the relative combinatorial inhibition, compared to non-treated cells, based on the whole-well viability assays, which was used in the prediction model for single agent total viability responses. The dark-blue parts of each bar indicate the expected additive inhibition from the combinations and the light-blue parts mark the unselective co-inhibition synergy based on the whole-well viability assays (excess inhibition% based on ZIP synergy model). **(b)** The relative inhibition of malignant AML cell subpopulations compared to inhibition of non-malignant cells (T cells and NK cells) in the patients based on the flow cytometry assay. **(c)** UMAP visualizations of the cell subpopulations based on the scRNA-seq transcriptomic profiles of the patient cells extracted before *ex vivo* compound testing. Cell clusters were identified using our ScType^31^ scRNA-seq processing pipeline with the Louvain clustering implemented in Seurat v3.1.0^32^. The identified clusters were annotated based on cell-specific marker information from our ScType marker database and unassigned cell types were manually identified (detailed in Method section). **(d)** The cell type composition of the patient samples based on their scRNA-seq transcriptomic profiles.

### Cellular heterogeneity of the patient samples explains variable treatment responses

We charted the cell type composition of each patient sample using the whole-transcriptome scRNA-seq profiling of the primary patient cells, extracted before the *ex vivo* compound testing (**Figure 3c-d**). Using our ScType^31^ scRNA-seq data processing pipeline and comprehensive cell marker database, which together with the other scRNA-seq analysis tools guarantee maximal specificity across both the cell clusters and cell types^32^, we observed a wide spectrum of cellular heterogeneity between and within the AML patient samples that provided further insights into the compound-induced cell subpopulation co-inhibition effects *ex vivo* and *in vivo*. For instance, the scRNA-seq transcriptomes identified NK cells from the diagnostic and refractory samples of AML3 patient case, therefore enabling the evaluation of co-inhibition effects of the patient-specific combinations on the NK cell population as well, in addition to T cells, with the combinatorial flow cytometry assay (**Fig. 3b**).

We also identified a number of potential molecular predictors of the treatment outcomes. For instance, the CD34+ blast cell clusters of AML1 and AML2 patient samples showed a high expression of two prognostic markers, ankyrin repeat domain 28 (*ANKRD28*) and guanine nucleotide binding protein 15 (*GNA15*) (**Suppl. Fig. 3**), which are associated with a significantly poorer overall survival in AML patients^33^. Specifically, in AML2 patient, our deconvolution approach identified a population of CD34+ blast cells that were further classified into two subpopulations (**Suppl. Fig. 4**); those expressing interferon-stimulated genes (ISGs), such as *ISG15, IFIT2* and *IFIT3* or not. Since the patient was not earlier treated with IFN-a/β^34^, and he did not have a known history of virus infection during the time of sampling^35^, it is likely that the ISGs+ CD34+ blast cells are drug resistant quiescent (G0) leukemic cells^36^, induced by induction therapy, and thereby causing the therapy failure, as indicated by change to azacitidine therapy after two cycles of induction therapy (**Suppl. Table 1**).

As each patient sample comprises different molecular backgrounds, we revealed patient-specific *ex vivo* synergies between multiple compounds. For example, venetoclax and losmapimod combination was validated to be synergistic in AML1 and AML2 patients, but not in the diagnosis and refractory samples of AML3 patient (**Fig. 2, Fig. 3**). The predicted and observed synergy and AML-selectivity of this combination may be partially explained by higher expression level of the primary target of losmapimod, *MAPK14* (**Suppl. Table 3**), both in the AML1 and AML2 patients’ AML cells in comparison to the two other samples (P<2.2×10^−16^, Wilcoxon rank-sum test), together with lower expression of *MAPK14* in non-malignant cells as compared to AML cells of both AML1 and AML2 patient samples (**Suppl. Fig. 5a**). We highlight below the most interesting combinations identified in each patient case separately, using our experimental-computational platform, together with their potential molecular determinants.

#### AML1 patient (diagnosis stage)

We first applied the integrated platform to predict patient-specific combinations for a 35-year old male AML patient (M2 subtype), with mutated *CEBPA, WT1 and CCND2*, and who was treated with induction chemotherapy (cytarabine-idarubicin, **Suppl. Table 1)**. Single-cell RNA-seq analysis showed that the patient sample was characterized by a high proportion of blasts (68%, **Suppl. Table 1**). The *ex vivo* single agent responses revealed high selective activity (sDSS>20) of crenolanib (FLT3 and PDGFR inhibitor), erlotinib (EGFR inhibitor), and ruxolitinib (JAK1&2 inhibitor) (**Suppl. Table 2**). In general, the patient sample showed highly selective sensitivity to multiple kinase inhibitors in addition to cytotoxic chemotherapy (**Suppl. Fig. 6**).

For combinatorial targeting of AML cells of AML1 patient cells, the predictive model identified increased synergy between molibresib (BET family inhibitor) and verdinexor (XPO1/CRM1 inhibitor) (**Fig. 3b**). This combination was unique to the AML1 patient sample (**Fig. 2c**), possibly because the leukemic cells in this sample expressed a higher level of the molibresib targets (*BRD2, BRD3* and *BRD4*, **Suppl. Table 3**), when compared to the non-leukemic cells of AML1 sample (P<0.0001, Wilcoxon rank-sum test, **Suppl. Fig. 5b-d)**. This unanticipated combinatorial effect would have remained unpredictable based on the single-agent testing results only.

#### AML2 patient (refractory stage)

The second patient case was a 68-year old male AML patient with a history of non-Hodgkin’s lymphoma and mutated *DNMT3A, ERG, U2AF1* and *BCOR* (**Suppl. Table 1**). Before sampling, the patient was treated with induction therapy (cytarabine-daunorubicin), followed by azacitidine, as the patient did not respond to the induction therapy (**Suppl. Fig. 7)**. The monocytic blast cell population of AML2 patient had the highest expression of cytidine deaminase (CDA), a known cytarabine inactivating enzyme that converts cytarabine to cytarabine-uracil^37^, possibly explaining the lack of the patient’s response to induction therapy (**Suppl. Fig. 5e)**. The single agent screening revealed sensitivity to epigenetic modifiers, apoptotic and heat shock protein inhibitors (**Suppl. Fig. 6**). Furthermore, the sample showed sensitivity to BAY87-2243 (mitochondrial complex I inhibitor), pevonedistat (NAE inhibitor) and plicamycin (RNA synthesis inhibitor), as the top monotherapies (**Suppl. Table 2**).

We identified a highly synergistic and low-toxic combination between ruboxistaurin (PKCβ inhibitor) and ipatasertib (AKT inhibitor), which was unique to the AML2 sample (**Fig. 3b**). This selective combination may appear since the non-leukemic cells of AML2 sample uniquely expressed the ruboxistaurin targets *PRKCB* and *PIM3* (**Suppl. Table 3**), as compared to the other samples that expressed neither of the targets in their AML cells (**Suppl. Fig. 5f-g**). Furthermore, the predictive approach revealed a strong patient-specific synergy among cyclin-dependent kinase (CDK) inhibitors and other single agents for this patient case. One such combination with a strong synergy was between AZD-5438 (a small-molecule CDK1/2/9 inhibitor) and abemaciclib (CDK4/6 inhibitor). This combination is interesting in the mutated *DNMT3A* AML2 patient samples, since simultaneous inhibition of CDK4 and CDK2 has shown to induce strong senescence response and growth arrest in leukemic cells with low DNMT3A expression^38,39^.

#### AML3 patient (diagnosis stage)

The final application case was to predict potential drug combinations for the diagnosis stage of 70-year old male AML3 patient (M1 subtype), with mutated *NPM1*, and previous history of Non-Hodgkin’s lymphoma (**Suppl. Table 1)**. In support of earlier observations that blast cells in AML patients with *NPM1*-mutation subtype are often CD34-negative^40^, the presence of CD34-leukemic cells was confirmed with our scRNA-seq analysis. Further analysis revealed a high proportion of erythroid lineage cells in the bone marrow mononuclear cells (35.8%), characterized by upregulation of hemoglobin related genes (e.g. *HBA2, HBB*), which could be the result of fetal hemoglobin induction due to azacitidine treatment, as is often observed in older AML patients^41^. The diagnostic stage *ex vivo* responses to the 456 single agents revealed navitoclax (Bcl-2/Bcl-xL inhibitor), dexamethasone and methylprednisolone (glucocorticoids), as the top 3 drugs with the highest activity in the patient sample (**Suppl. Table 2**). The potency of navitoclax is in line with our earlier observation of higher activity of Bcl-2 inhibitors in M1 subtype AML^42^. In general, the patient sample showed a high selective sensitivity (sDSS range 13-21) to a number of targeted compounds, including kinase inhibitors, HSP inhibitors and apoptotic modulators (**Suppl. Fig. 6**).

The predictive platform identified a strong and unique synergy between patupilone (microtubule inhibitor) and uprosertib (AKT kinase inhibitor) in the AML3 patient diagnostic sample (**Fig. 2c**). It has been shown that a dual inhibition of microtubules and AKT/mTOR pathways can synergistically enhance apoptosis of cancerous cells, particularly in cancer patients resistant to mitotic inhibitors, since PI3K/Akt/mTOR pathway is associated with resistance in multiple cancers including leukemia^43,44^. In line with this observation, patupilone primary target, tubulin beta (TUBB, **Suppl. Table 3**), showed higher expression in the leukemic cells of AML3 patient diagnosis sample, compared to non-leukemic cells (P<0.0001, Wilcoxon rank-sum test, **Suppl. Fig. 5h**), suggesting that the synergistic co-inhibition might be achieved through selective inhibition of the tubulins involved in cancer chemoresistance and metastasis^45,46^.

#### AML3 patient (refractory stage)

The leukemia of patient AML3 became refractory to azacytidine, and the patient was therefore transferred to a venetoclax regimen after two months. The *ex vivo* single agent responses revealed that the refractory stage sample was still sensitive to dexamethasone (glucocorticoid), BMS-754807 and GSK-1904529A (IGF1R inhibitor) (**Suppl. Table 2**), similar to the diagnostic sample (**Suppl. Fig. 6**). However, the refractory sample became more sensitive both to epigenetic modifiers (P=1.5×10^−3^) and chemotherapies (P = 1.2×10^−3^, Wilcoxon rank sum test).

Our scRNA-seq analysis revealed a decreased level of erythroid like cells, together with the loss of CD16+ monocytes and mast cell population in the refractory stage of AML3 patient, as compared to the diagnosis stage (**Fig. 3d**). Interestingly, the synergy between ruboxistaurin (PKCβ inhibitor) and ipatasertib (AKT inhibitor), which was observed only in the diagnostic sample, was lost as the disease progressed from diagnosis to refractory stage, possibly due to the lower expression ruboxistaurin-target *PIM3* in the leukemic cells of the refractory sample (**Suppl Fig. 4g**). This case example demonstrates the need for longitudinal patient sampling and subpopulation-level prediction machinery to dynamically tailor AML-selective combinations at different disease stages.

## Discussion

AML cells harbor a number of genetically and epigenetically heterogeneous subpopulations, each of which may be associated with a distinct cellular function or phenotype such as self-renewal, drug sensitivity or resistance. To address this heterogeneity in personalized treatment design, we implemented a computational-experimental platform and applied it to AML patient primary cells to rationally combine drugs that preemptively inhibit multiple AML-related dysfunctions or resistance mechanisms in individual patients. The selective approach avoids a severe co-inhibition of non-malignant cells, with the aim to improve both combination efficacy and tolerability. To the best of our knowledge, this is the first systematic approach to personalized drug combinations selection that takes into account both the molecular heterogeneity of AML cells and the possible toxic effects of combinations, thereby increasing their likelihood for clinical translation. The approach requires only a limited number of patient primary cells, and it is widely applicable to hematological cancers that are a tractable disease system amenable to scRNA-seq profiling and systematic *ex vivo* compound testing. In the present work, we used a total of 30-40 million cells per patient both for the primary single agent screening and scRNA-seq profiling, as well as for the combinatorial validation experiments with whole-well and flow-based assays (a vial of 10 million cells for each assay). The implementation of the experimental-computational approach takes less than three weeks per sample, where scRNA-seq is the current bottleneck, demonstrating the approach can be applied within a clinically actionable timeframe, especially once the sequencing becomes faster.

Beyond AML, there is an urgent clinical need to develop rational and systematic strategies for designing anticancer combinatorial treatment that can target cancer heterogeneity and drug-resistant cell populations^47^. Earlier approaches have used experimental testing of single drug efficacies in genetically variant cell subpopulations, together with RNA interference to model the functional diversity and perturbation effects of heterogeneous tumors, followed by computational optimization algorithm that predicts how drug combinations will affect heterogeneous tumors^48^. While such optimized design principles can offer useful insights into intratumoral heterogeneity, profiling each component subpopulations with single drugs together with knock-down experiments in patient cells is arguably not amenable to routine clinical application. Recently, single-cell RNA-sequencing has been widely-utilized to chart the cellular composition of complex samples, and to design combinatorial regimens in solid tumors that show increased treatment efficacy over monotherapy^49^. Even though scRNA-seq enables efficient mapping of primary AML cell populations at baseline^23^, when used alone, it lacks the possibility to profile functional responses of drug treatments. Another recent study utilized nested effects modeling combined with Mass Cytometry Time-of-Flight (CyTOF) to simultaneously profile single agent responses and to catalogue signaling heterogeneity at the single-cell level, with the aim to suggest drug combinations that lead to maximal desired intracellular effects^50^. However, this method requires CyTOF profiling before and after the drug treatment, hence limiting its predictive clinical value.

Next-generation functional precision medicine approaches that combine bulk molecular and genomic characterization of individual patient samples at various disease stages with the whole-well ex-vivo phenotypic profiling of the patient samples using large compound libraries has demonstrated to provide a relatively fast and practical platform for translational discoveries in hematological malignancies both for monotherapies and combination discoveries^51-54^. High-throughput flow cytometry-based profiling approaches have been stablished to further evaluate the *ex vivo* sensitivity of various cell populations in primary AML samples^42^, where especially refractory patient samples often have lower blast cell percentages, making the whole-well viability assays less reliable for drug response profiling. However, flow cytometry profiling of a larger number of drug combinations in multiple doses is currently not feasible in scarce primary patient cells; instead, translational strategies rely on testing only selected drugs or combinations in single concentrations^55^. Therefore, there is a need for efficient approaches to prioritize the most potent and selective combinations for the subpopulation-level and further preclinical validation. Our experimental results demonstrated that the bulk viability screening assay captures surprisingly high predictive power for the AML cell subpopulation co-inhibition effects, when combined with the scRNA-seq transcriptomic profiling and machine learning modelling. The whole well synergy assay should therefore be considered as the first experimental filtering phase, following the model predictions, which enrich a smaller number of the most synergistic and selective combinations to be validated in the flow-based subpopulation assays, before entering the clinical translation phase.

The benefits of the integrated approach were demonstrated in multiple case examples, including the combination between saracatinib (SRC, ABL inhibitor) and cytarabine (DNA synthesis inhibitor), which was confirmed to be highly synergistic, low-toxic and selective for AML1 patient (**Fig. 2 and 3**). The combination is likely to remain effective also in more complex disease systems, since the SRC family kinases are activated in AML stem/progenitor cells, known to contribute to their survival and proliferation^56^. Furthermore, an *in vivo* study has shown that SRC kinase inhibitor treatment enhances chemotherapy-induced targeting of primary murine AML stem cells, capable of regenerating leukemia in secondary recipients by activating of p53,^57^ and therefore this combination may be best suited for cancers that contain wild-type p53, as is the case with AML1 patient (**Suppl. Table 1**). On the other hand, a cytarabine-saracatinib combination showed synergy in the whole-well viability assay only in one of the two predicted samples (**Fig. 2a**), indicating that part of the heterogeneity in the combinatorial response is either not captured by the whole-well viability readouts or remains unpredictable by the integrated approach. We also demonstrated that combinations having high synergy in the whole-well viability assay may sometimes lead to combinations with high likelihood of toxic effects, especially in patient samples with limited amounts of non-leukemic cells, making the validation experiments using subpopulation-specific response assays highly informative for translational applications. Notably, the flow-based experiments revealed a wide inter-patient variability in the combinations involving venetoclax. For instance, the combination of venetoclax and losmapimod showed a relatively variable *ex-vivo* synergy-efficacy-toxicity profile between diagnostic AML1 and refractory AML2 samples (**Fig. 3**).

Given that drug combination synergy is a rare phenomenon, with estimated likelihood of synergy being around 5% in large-scale combination screens^19,58,59^, our predictive platform led to highly accurate combination prioritization, achieving a higher than 50% success rate in the validation experiments. For specific patient cases, the success rate of the predictions was even higher. For instance, in the AML1 patient, we tested 17 of the predicted combinations using the whole-well viability assay, and the experimental validations revealed that 10 (59%) of the combinations showed strong synergistic effect (ZIP>5), and 15 (88%) of the predicted combination were either synergistic or additive (ZIP>0). We acknowledge that some of the *ex vivo* predictions may not lead to clinically actionable therapies. For instance, simultaneous blocking of both topoisomerase I and II showed synergies in all the patient samples. While this combination might provide an opportunity to effectively co-target AML rescue pathways, as inhibition of either of the enzymes alone may lead to increase in the expression of the other, hence resulting in treatment of resistance^24^, the sequential or simultaneous co-inhibition of topoisomerase I and II by compounds forming DNA lesions has shown severe side effects^60^. Therefore, dual inhibitors with a longer half-life and improved binding efficacy (e.g. tafluposide and batracylin) are being designed to tackle the treatment resistance and toxicity^24^. Similarly, the combination of AZD-5438 (CDK1/2 inhibitor) and abemaciclib (CDK4/6 inhibitor) showed patient AML2-specific synergy; however, this combination is unlikely to be translated to clinics, as AZD-5438 monotherapy trial was discontinued due to intolerable toxicities when administered continuously in patients with advanced solid tumours^38^.

These AML case studies also demonstrated that in addition to the overall co-inhibition synergy, it is important to also consider the relative differences between the combination efficacy and toxicity, since these are critical determinants for the clinical success of a therapy^11,18,61^. As a unique component of our machine learning model, it makes use of the information from scRNA-seq on the target expression levels of the compounds both in malignant and non-malignant cells, when predicting not only synergistic effects but also the differential co-inhibition effects between malignant and non-malignant cells. To the best of our knowledge, there are no large-scale studies of such differential co-inhibition effects in cancer cells, but it is arguably even a rarer phenomenon than an overall combination synergy alone. Notably, the flow assay validations showed that 67% of the predicted and synergistic combinations had highly selective co-inhibition effect (<50% toxicity to T and NK cells), indicating that the predictive approach effectively avoids broadly toxic and potentially synergistic combinations that unselectively kill various cell types. Combination synergy, efficacy and toxicity profiling each provide complementary information on the combinations’ mode-of-action, and should therefore be informative for translational applications. Our machine learning model enabled also possibilities to identify potential molecular markers for the low-toxic and synergistic combinations in each patient case, based on their scRNA-seq profiles, including *MAPK14* for venetoclax and losmapimod combination, which warrant further testing as predictive biomarkers in larger AML patient cohorts, toward safe and effective combination therapies.

## Materials and Methods

### Patient samples

Four bone marrow aspirates from three AML patients (indexed as AML1, AML2 and AML3, the latter with both diagnostic and refractory samples) were collected at Helsinki University Hospital and The Finnish Hematology Registry and Clinical Biobank (FHRB) after informed consent, using protocols approved by a local institutional review board in accordance with the Declaration of Helsinki. Patient clinical characteristics and mutation profiles are shown in **Suppl. Table 1.**

### Single-cell RNA sequencing

Enriched (Ficoll-Paque PREMIUM; GE Healthcare) mononuclear cells were used for Chromium Single Cell 3’RNA-sequencing. The gel beads in emulsion (GEM) generation, cDNA amplification and library preparation were performed according to the manufacturer’s instructions using Chromium Single Cell 3’ v2 Reagent Kit (10x Genomics) aiming for a 4000-6000 target cell capture per sample. The sample libraries were sequenced in Illumina HiSeq 2500 using Rapid mode aiming for sequencing depth of 50 000 reads per cell. The Cell Ranger v1.3 mkfastq and count analysis pipelines (10x Genomics) were used to demultiplex and convert Chromium single cell 3’ RNA-sequencing barcode and read data to FASTQ files and to generate align reads and gene-cell matrices. The gene-barcode matrices were analyzed using the ScType^31^ scRNA-seq processing pipeline with the Louvain cell clustering as implemented in Seurat v3.1.0^32^. Cells with either a low (below 100-200) or unexpectedly high number (3000-4000, depending on the sample) of detected genes, or cells with more than 8-10% of mitochondrial UMI counts, were classified as low-quality or uninteresting cells and were excluded from the analysis.

### Cell type identification and manual annotation

Cell-specific marker information was automatically extracted from our ScType marker database^31^, and unassigned cell types were manually identified based on the specific expression markers:

| Cell types | Markers |
| --- | --- |
| Monocytes | CD14+ |
| T cells | CD3+, CD4+,<br>CD8+ |
| Macrophages | APOC1+ CD71+ |
| Erythroid-like and erythroid precursor cells | GYPA+, HBB+ |
| Granulocytes | MPO+ |
| Blasts | CD34+, CD33+<br>CD45<sup>low</sup> |
| Natural killer cells | CD56+ GZMB+,<br>CD335+ GZMB+ |
| Neutrophils | MPO+ AZU1+ |
| Mast cells | FCER1A+<br>CD203c+ |

### Compound sensitivity testing

We used a library consisting of 456 single compounds from the FIMM oncology set, which is a comprehensive and evolving compound collection of both approved drugs and investigational compounds (**Suppl. Fig.1**). For the single agent response testing^51^, 20 μL of fresh AML cells (approx. 10 000) suspension in Mononuclear Cell Medium was added per well to pre-drugged plates with 10-fold dilution series of five concentrations, and the whole-well cell viability was measured with CellTiter-Glo (CTG, Promega) in duplicate, as previously described^19,51^. The concentration ranges were selected for each compound separately to investigate their full dynamic range of the dose-response relationships. After 72 h of incubation at 37°C and 5% CO_2_, cell viability of each well was measured using the CTG luminescent assay and a Pherastar FS (BMG Labtech) plate reader. The percentage inhibition was calculated by normalizing the cell viability to negative control wells containing only 0.1% DMSO and positive control wells containing 100 μM cell killing benzethonium chloride (BzCl).

The dose-response curve of each drug was fitted using the four-parameter nonlinear logistic regression, also called the Hill equation:

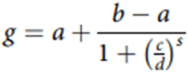

where *g* is a response of single agent at dose *d, a* is the minimum asymptote, *b* is the maximum asymptote, *c* is the half-maximal effective concentration (relative IC_50_) and *s* is the slope of the curve. The fitting was done using the drc package (version, 3.0-1) in R. The maximal relative inhibition level closest to relative IC_50_ concentration was used as single agent response outcome measure when training the machine learning model **(Suppl. Fig. 2a**). We quantified the summary response of each compound in the patient samples using the drug sensitivity score (DSS), as described previously^62^. Selective DSS (sDSS) was calculated by subtracting from the patient-specific DSS the average of 6 healthy bone marrow samples’ DSS values (**Suppl. Table 2**).

### Compound-target information

We retrieved comprehensive compound-target information from two publicly available databases: DrugTargetCommons v2.0 (drugtargetcommons.fimm.fi)^63,64^ and DGIdb v 3.0 (www.dgidb.org)^65^. We included in the compound-target networks only those protein targets with a relatively potent binding affinity (K_d_, K_i_, IC_50_<1000nM) from Drug Target Commons database and protein targets with a reference score > 2 in DGIdb database. Using this rather liberal potency cut-off, we retrieved 898 unique protein targets for the 456 compounds, including both their nominal on-targets as well as potent off-targets. The predictive modelling made use of the comprehensive compound-target interaction networks, ranging from primary targets to downstream targets and other modifiers of the compounds’ responses to make systematic predictions of their co-activities based on a wide spectrum of binding affinities and inhibitory levels of the compounds against target pathways. However, when interpreting the identified co-inhibition effects of the combinations in specific patient samples, we only focused on primary targets and off-targets with similar affinity or inhibition than the intended targets, using target information from DrugBank (https://www.drugbank.ca/)^66^ IUPHAR^67^, and MedChemExpress (https://www.medchemexpress.com/) (**Suppl. Table 3**).

### Predictive modelling

Since systematic testing of all the potential combinations among the 456 compounds is not experimentally feasible (>100,000 pairwise combinations), we adopted an efficient computational strategy that combines the multiple compound-target expression levels from scRNA-seq data into a single compound-specific target score for each individual cell using the gene set variational analysis (GSVA) approach^21^. More specifically, we reduced the dimensionality of the genome-wide scRNA-seq transcriptomic profiles and calculated the target enrichment score for individual cells using GSVA, which calculates the normalized difference in empirical cumulative distribution functions of gene expression ranks inside and outside the drug proteins targets (here, gene set), as an enrichment statistic per cell, which is further normalized by the range of the statistic values across all the gene sets. These compound-target scores were used as input for training a regularized boosted regression trees (XGBoost) algorithm to predict the single agent viability inhibition (measured as response nearest to the relative IC_50_ concentration, see **Suppl. Fig. 2a**) for each compound and each patient separately. The predictive model was optimized using Bayesian optimization with repeated 5-fold cross-validation (CV) approach within the 456 compounds in each patient sample separately, and the optimized model was then used to predict the compound combination responses in the particular patient sample. The union of targets of individual compounds was used to construct a combination target-score, similar to our previous work,^19^ and this combination score was used as input for the trained patient-specific XGBoost model for the drug combination response prediction. ZIP synergy model^22^ was utilized to identify the most synergistic predicted combinations for each patient. In order to eliminate low confidence predictions, we utilized a conformal prediction (CP) approach^21^. In short, CP uses absolute values of the repeated CV residuals as dependent variable for the error model to predict how unlikely (conformal) each prediction is, which sub-clusters the prediction space into various confidence level regions. Specifically, we removed all the XGBoost predictions with a nonconformity score lower than 0.8. In doing so, we used in combination predictions only those single agents whose response could be predicted confidently using their target expression levels.

In order to prioritize the most promising and clinically actionable combinations for each patient individually, the predicted combinations were further ranked based on their predicted toxic effect on non-malignant cells, along with their predicted capacity to target multiple cancer cell subpopulations. First, we selected the top 10% combination based on the predicted ZIP synergy score. Out of the selected synergistic combinations, we further selected the top 10% combinations with the highest predicted efficacy on the malignant cells. Finally, we prioritized the top 10% of the remaining combinations with the lowest target enrichment scores in the non-malignant cells (i.e., lowest number of cells with positive GSVA values), thus excluding those synergistic and effective combinations that could potentially result in toxic effects. The underlying principle is that the model attempts to select those compounds and combinations, which follow our assumption of their mode-of-action. Namely, whenever a compound’s target gene set is highly enriched across all cell types, it is expected to have cell type-independent inhibition effects, which should lead to high CTG *ex vivo* responses. In contrast, the compounds with low target gene set enrichment in all cell types are assumed to show lower CTG *ex vivo* responses. We note that not all the compounds follow this assumption, and those compounds as well as the others that the model is not able to learn properly during Bayesian optimization, are removed with the conformal learning. Ultimately, the model is used to predict which combinations will produce the highest cell killing effect, and then we select among the top predictions those with a lower likelihood of toxic effects (e.g. de-prioritize combinations whose target gene set is highly enriched in healthy cell types).

### Validation of the predicted combination using CellTiter-Glo (CTG) viability assay

The predicted combinations involving 64 single agents were subsequently tested on the bone marrow mononuclear cells of each patient in 8×8 dose-response matrix using CTG viability assay, similarly as described before^19,51^ and above (see section Compound sensitivity testing). The combination synergy in the experimental validations was quantified using the zero interaction potency (ZIP) model22, calculated based on the dose region around IC_50_ values of each compound in combination, separately for each patient and combination, since the machine learning model predicted synergistic inhibition at doses closest to IC_50_ concentration.

### Investigation of the combination using high-throughput flow cytometry (HTFC) assay

We performed HTFC assays to assess the effects of drugs on multiple cell populations in primary AML patient samples, as described previosly^42^. Briefly, the compounds were dissolved in 100% DMSO or an aqueous solution and dispensed on tissue culture grade Corning V-bottom 96-well plates using an Echo 550 Liquid Handler (Labcyte) in 4 concentrations of 10-fold dilution series around the IC50 (**Suppl. Fig. 2b)**. The conditioned medium suspended cells were seeded at 100 000 cells/well on the drug plates and incubated for 3 days at 37°C and 5% CO_2_. The cells were washed with a cell staining buffer (PBS with 2% fetal bovine serum), centrifuged at 600×*g* for 5 min. To profile cell subpopulation responses, the cells were stained with BV605 Mouse Anti-Human CD56, BV421 Mouse Anti-Human CD19, BV421 Mouse Anti-Human CD3, FITC Mouse Anti-Human CD45, APC Mouse Anti-Human CD34, BV786 Mouse Anti-Human CD38 antibodies (all antibodies from BD Biosciences) for 30 min at room temperature in the dark. After antibody staining, the cells were washed in cell staining buffer and resuspended in 25 μL Annexin V binding buffer containing 0.5 μL of PE Annexin V and 7-aminoactinomycin D (7AAD) and incubated for 10 min at room temperature in the dark. The cells were analyzed on an iQUE Plus instrument (Intellicyt). The data were analyzed by using FlowJo software v10.6.2 (Treestar) and collected 50 to 80 000 events. Briefly, cell singlets were identified based on FSC-A vs FSC-H ratio and live cells were identified using Annexin V and 7AAD markers followed by identification of leukocytes based on CD45 and FSC-A. We further characterized NK cells (CD56 positive and CD3/CD19 negative), leukemic stem cells (CD34 and CD38 positive) and T/B cells (based on SSCA and CD3/19) from the leukocytes. The details of the gating strategy are illustrated in the **Suppl. Fig. 8.**

## Acknowledgements

The authors are grateful to the patients who have donated samples to the study. We thank the FIMM High Throughput Biomedicine and Single Cell Analytics units supported by HiLIFE and Biocenter Finland for their technical support, Disha Malani (FIMM) for information about the patient samples, Olli Kallioniemi (Science for Life Laboratory, Sweden) for financial support for the molecular and compound profiling, and Shobha Potluri (Senior Principal Scientist at Pfizer Rinat Laboratories, SF, USA) for support for the scRNA-seq data generation. The work was partially funded by the Academy of Finland (grants 310507, 313267, 326238 to TA; iCAN Digital Precision Cancer Medicine Flagship grant 1320185 to TA, CH), Cancer Society of Finland (TA, KW, CH), Sigrid Jusélius Foundation (KP, TA and KW), Business Finland (KP), Novo Nordisk Foundation (to KW; NNF17CC0027852), and Finnish Medical Foundation, Finnish Cancer Institute (MK).

## Supplementary Material

**Supplementary Figure 1.**
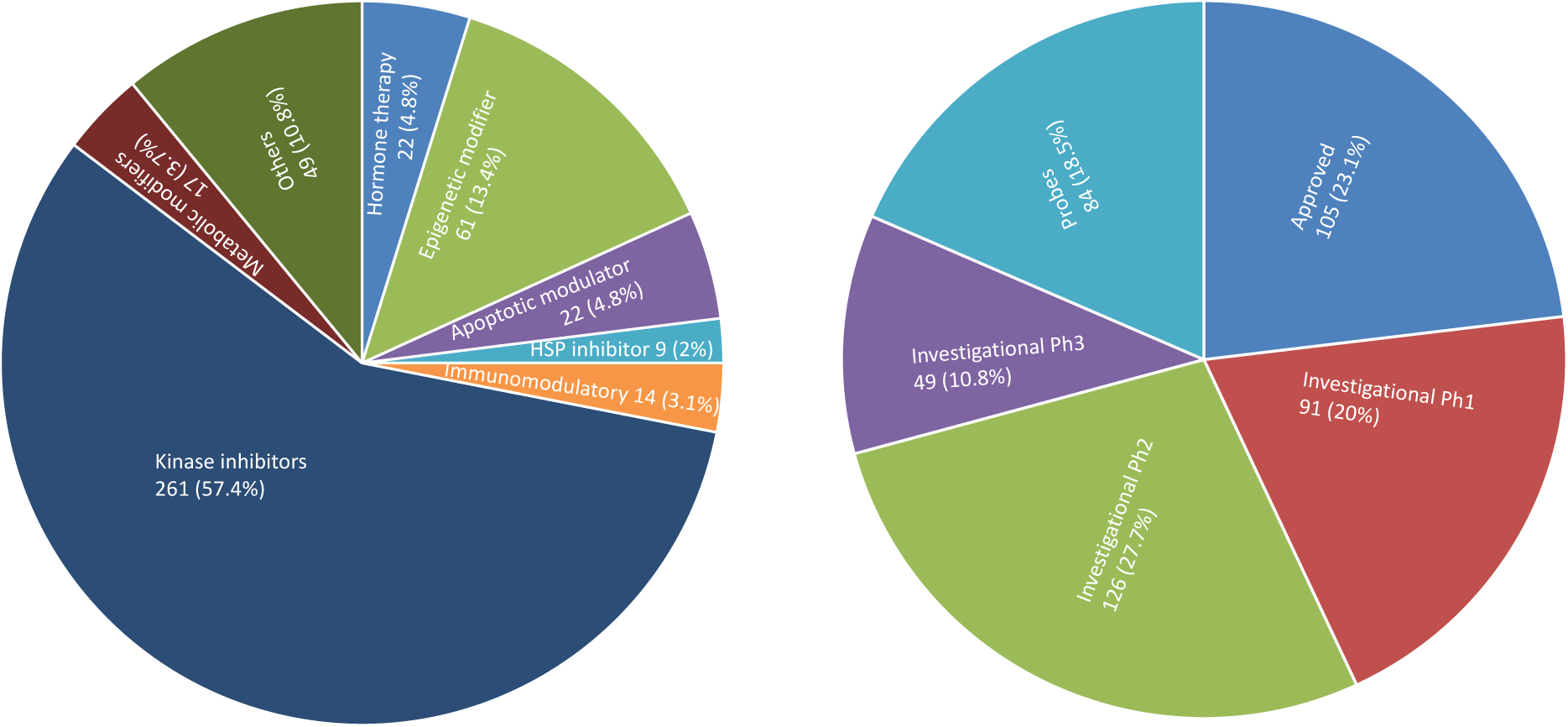
The mechanism of action (left panel), and the clinical development phase (right panel) of the 456 compounds used in the study.

**Supplementary Figure 2.**
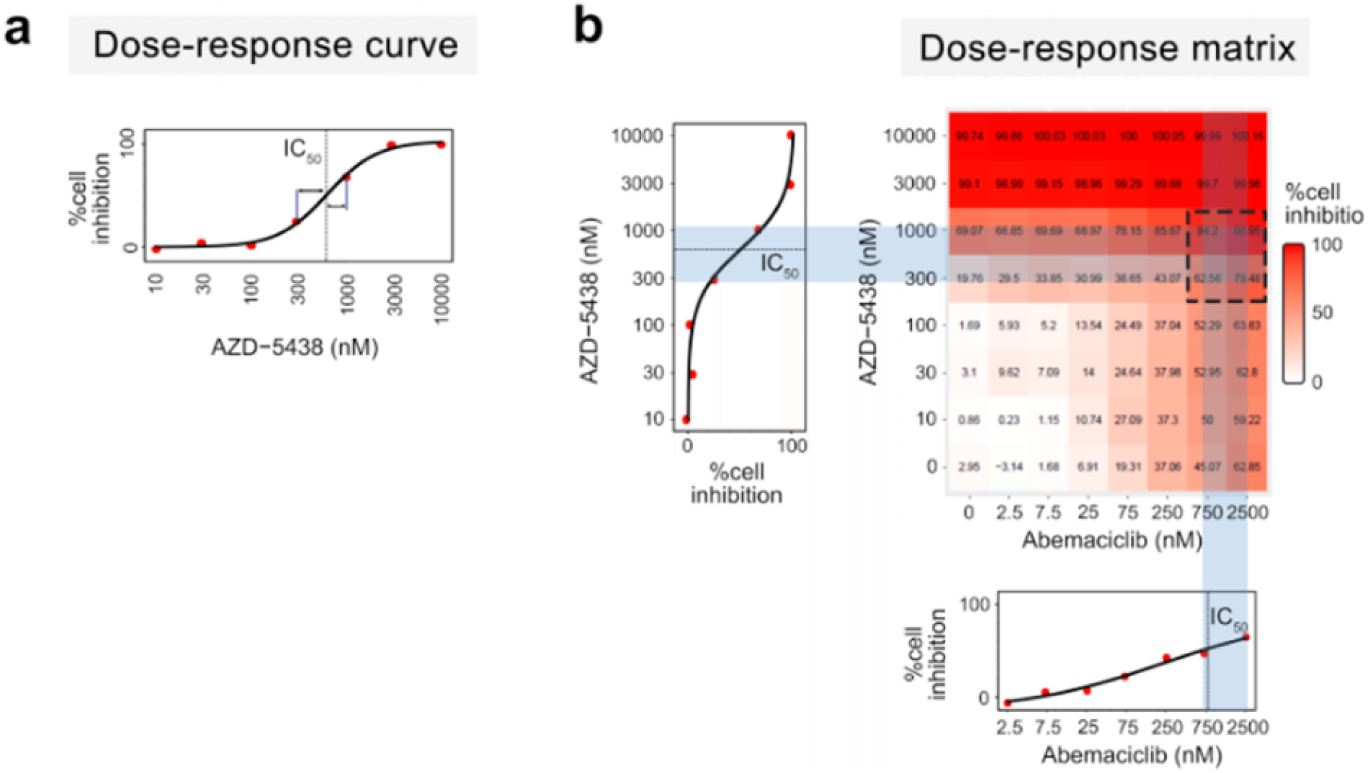
**(a)** An example of the compound dose selection utilized for calculating the compound inhibition level in the machine learning model training using single agent responses. The response at the dose closest to IC_50_ value (dashed line) in the logarithmic space is selected as the model input (cell inhibition percentage at 1000 nM in this example). (**b)** An example of drug combination dose selection used in flow cytometry assay validation. The overall cell viability inhibition at the two doses nearest to the relative half-maximal inhibitory concentration (IC_50_) of each single agent (bold dashed rectangle) was measured in the flow cytometry experiments.

**Supplementary Figure 3.**
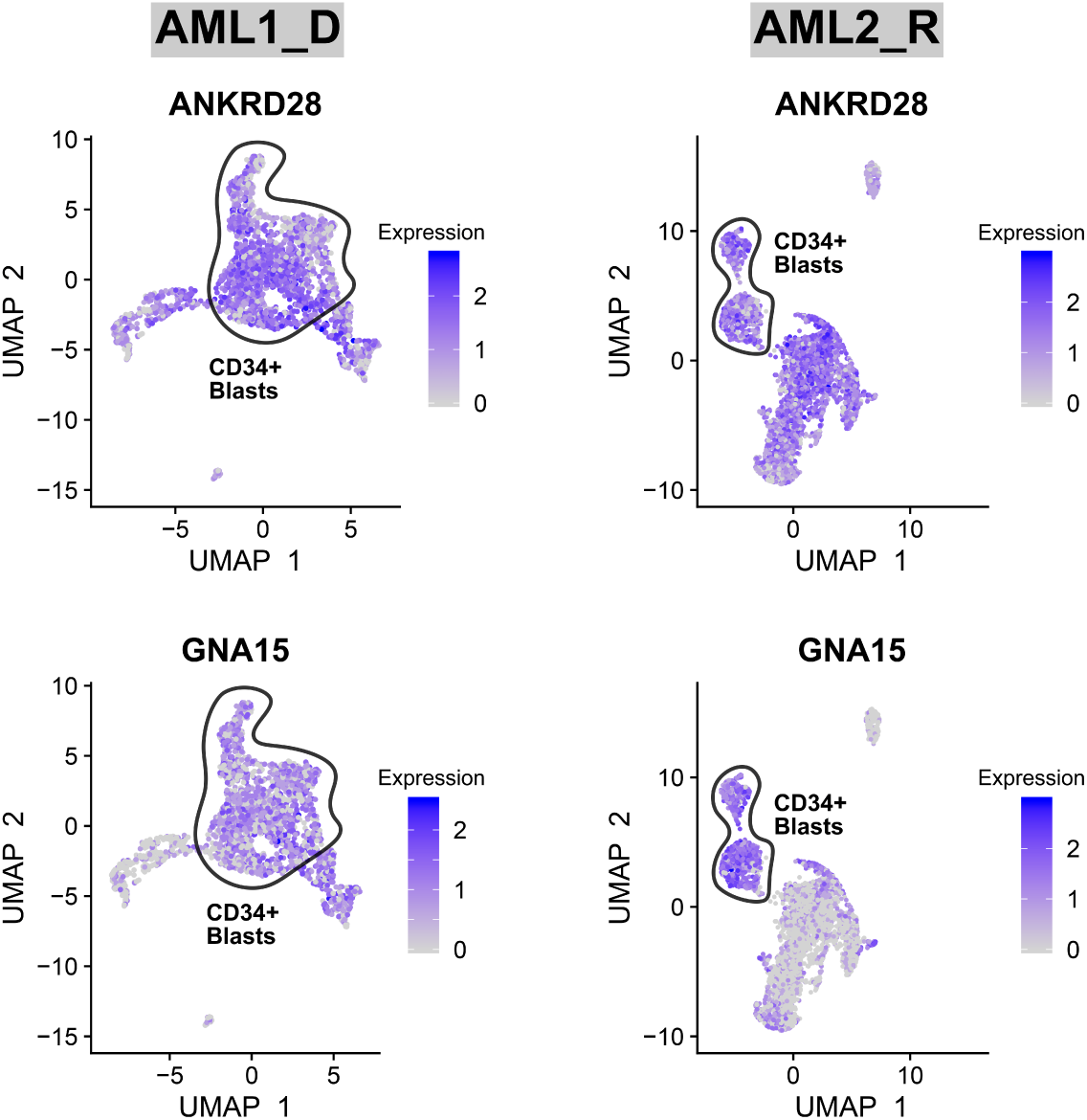
Expression of ANKRD28 and GNA15 markers, associated with a significantly poorer overall survival in AML patients, in the CD34+ blast cells AML1 and AML2 patient samples.

**Supplementary Figure 4.**
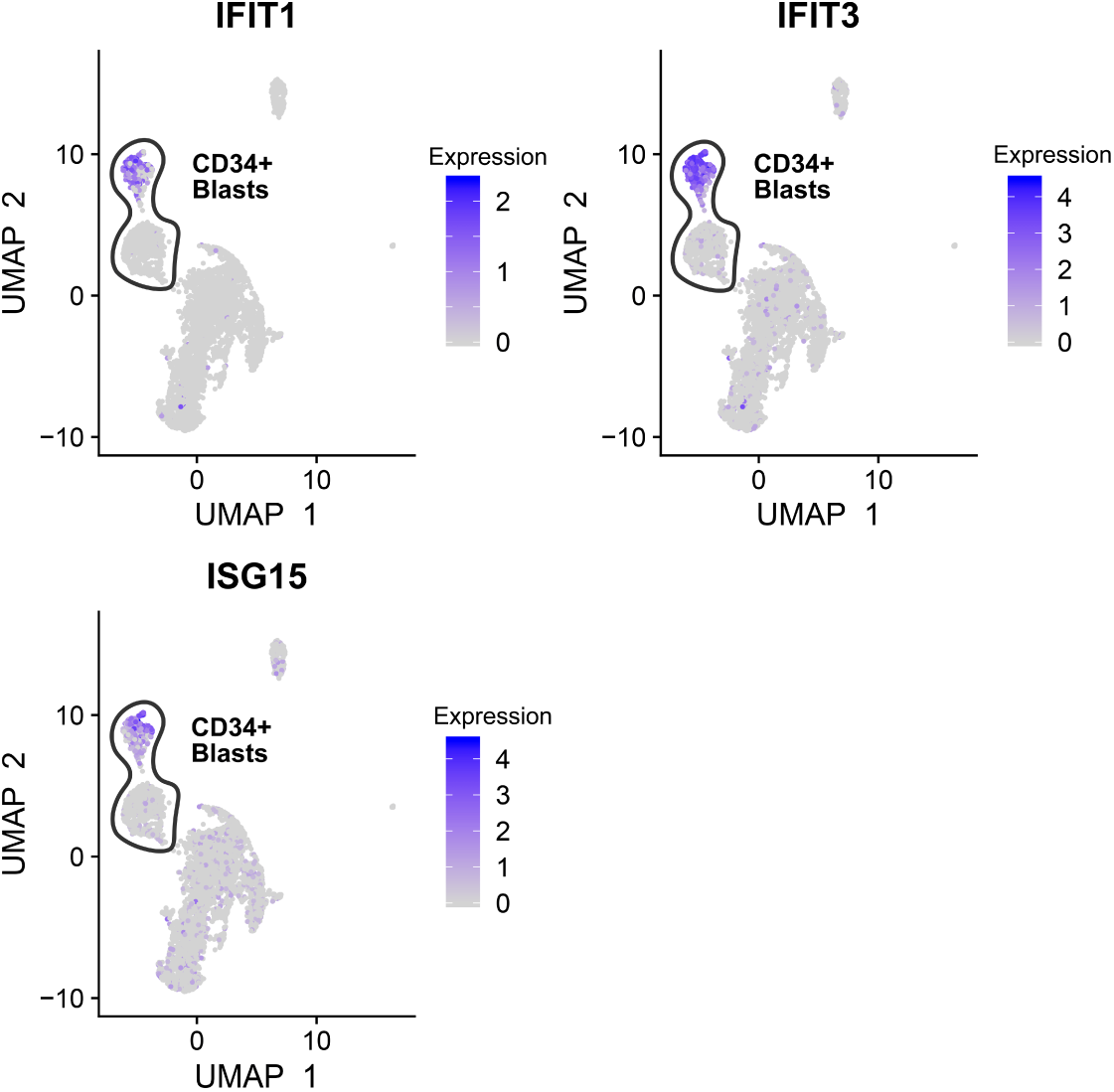
Expression of interferon-induced genes (IFIT1, IFIT3, ISG15) in the CD34+ blast cell population of the AML2 patient sample. The color represents the log normalized expression level of the genes obtained after sctransform of RNA count data.

**Supplementary Figure 5.**
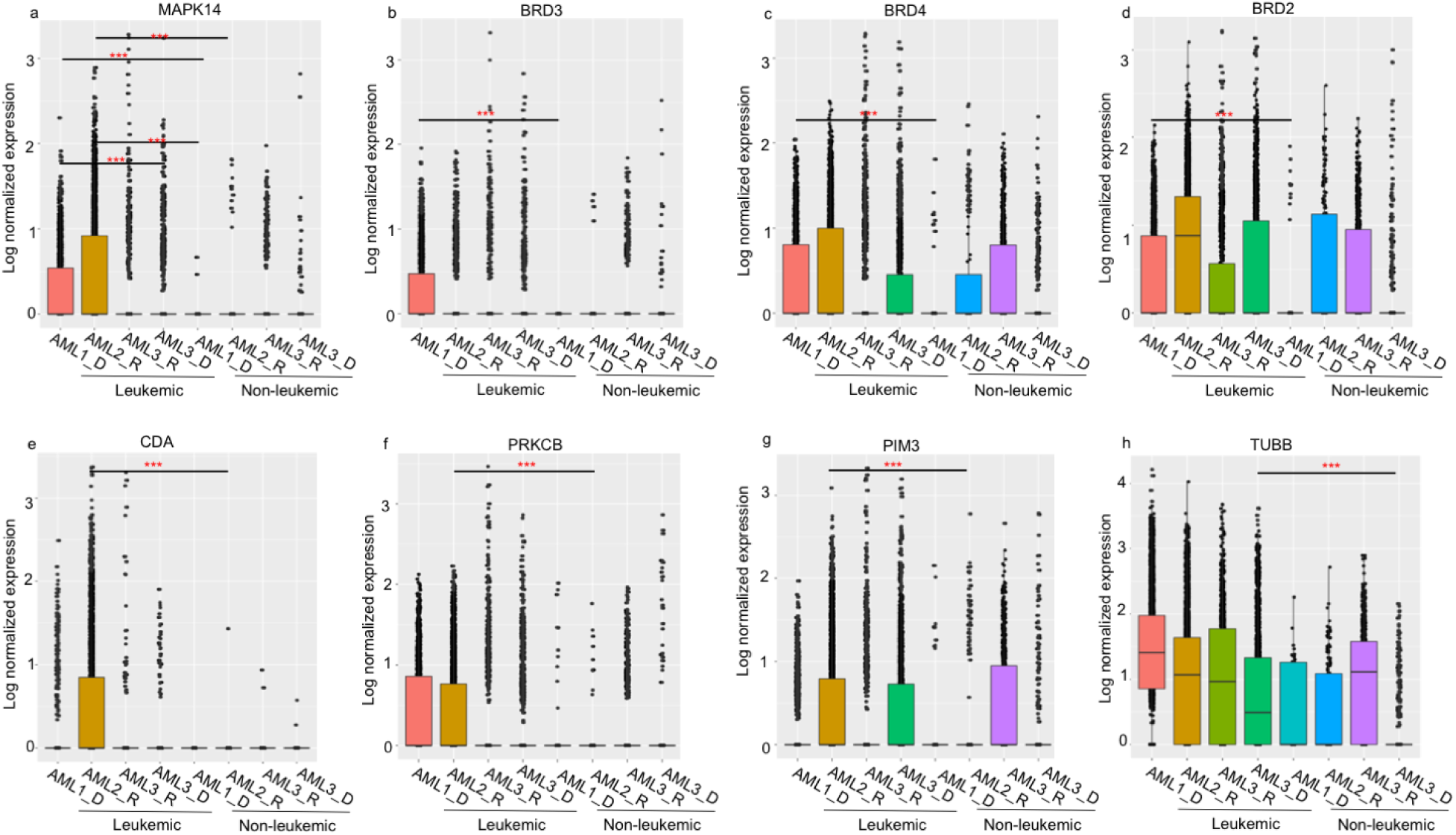
Target expression levels of compounds identified as effective and less toxic combinations in the AML samples. (**a**) Comparison of the expression of losmapimod primary target MAPK14 in the leukemic cells of AML1 and AML2 patients, against leukemic cells of AML3 patient, and non-leukemic cells of AML1 and AML2. (**b-d**) Comparison of expression levels of molibresib targets BRD2, BRD3, and BRD4 between leukemic and non-leukemic cells of AML1. (**e**) Comparison of the cytarabine deactivating enzyme cytidine deaminase (CDA) expression in the leukemic cells of AML2 patient with leukemic cells of other samples and non-leukemic cells of AML2. (**f-g**) Comparison of the expression levels of ruboxistaurin targets PRKCB and PIM3 in the leukemic cells of AML2 patient with leukemic cells of other samples and non-leukemic cells of AML2. (**h**) Comparison of expression levels of patupilone target tubulin B (TUBB) in the leukemic cells of AML3 patient with leukemic cells of other samples and non-leukemic cells of AML3. In each panel, y-axis shows log-normalized count values after sctransform^32^, where p-values were calculated using Wilcoxon rank-sum test. ***p<0.001.

**Suppl Figure 6.**
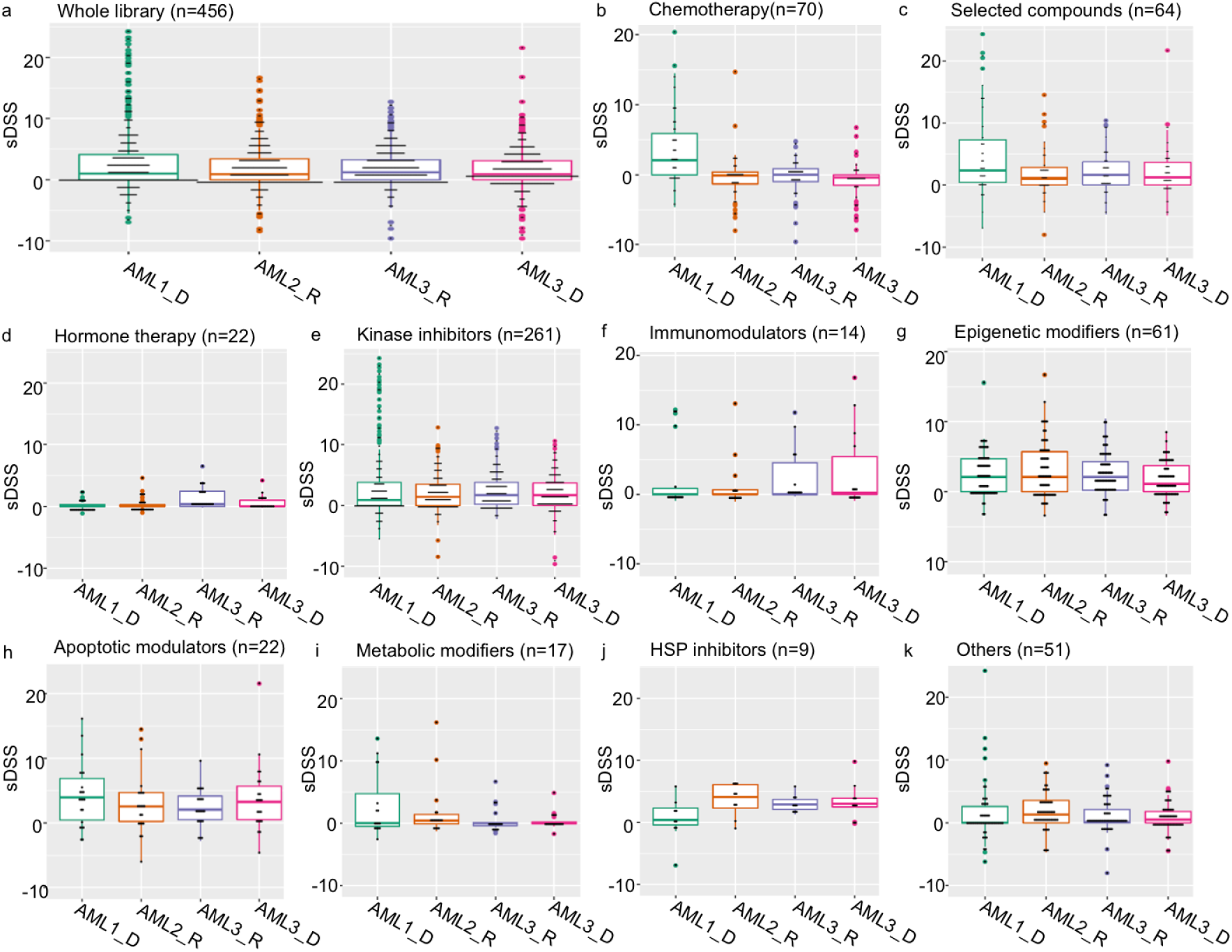
Comparison of *ex vivo* whole-well single agent sensitivity profiles across patient samples and compounds classes. Selective DSS (sDSS) was calculated by subtracting from the patient DSS the average of 6 healthy bone marrow sample DSSs.

**Supplementary Figure 7.**
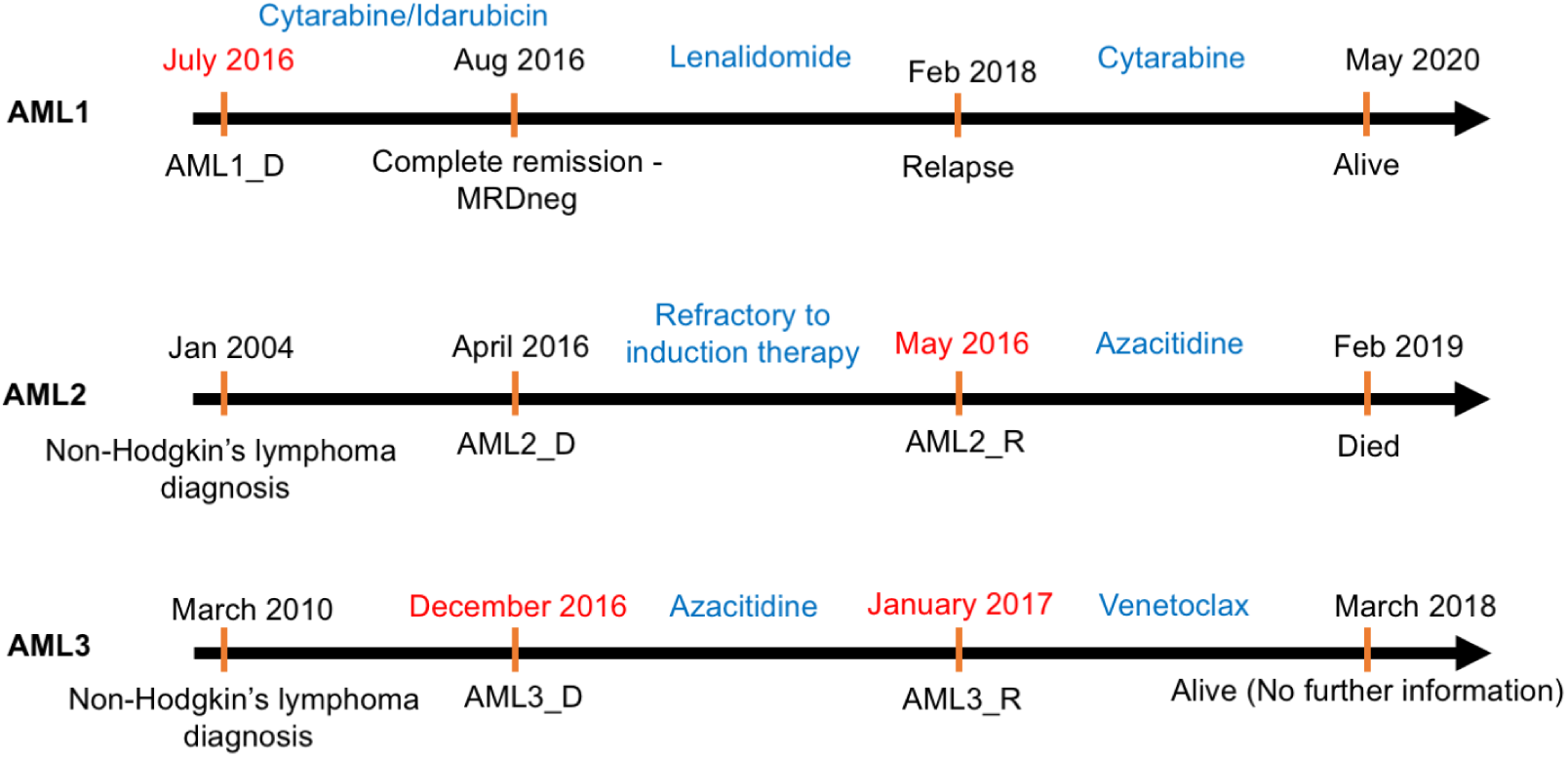
Timeline of the treatment regimens for the individuals around the time of sampling (marked in red font).

**Supplementary Figure 8.**
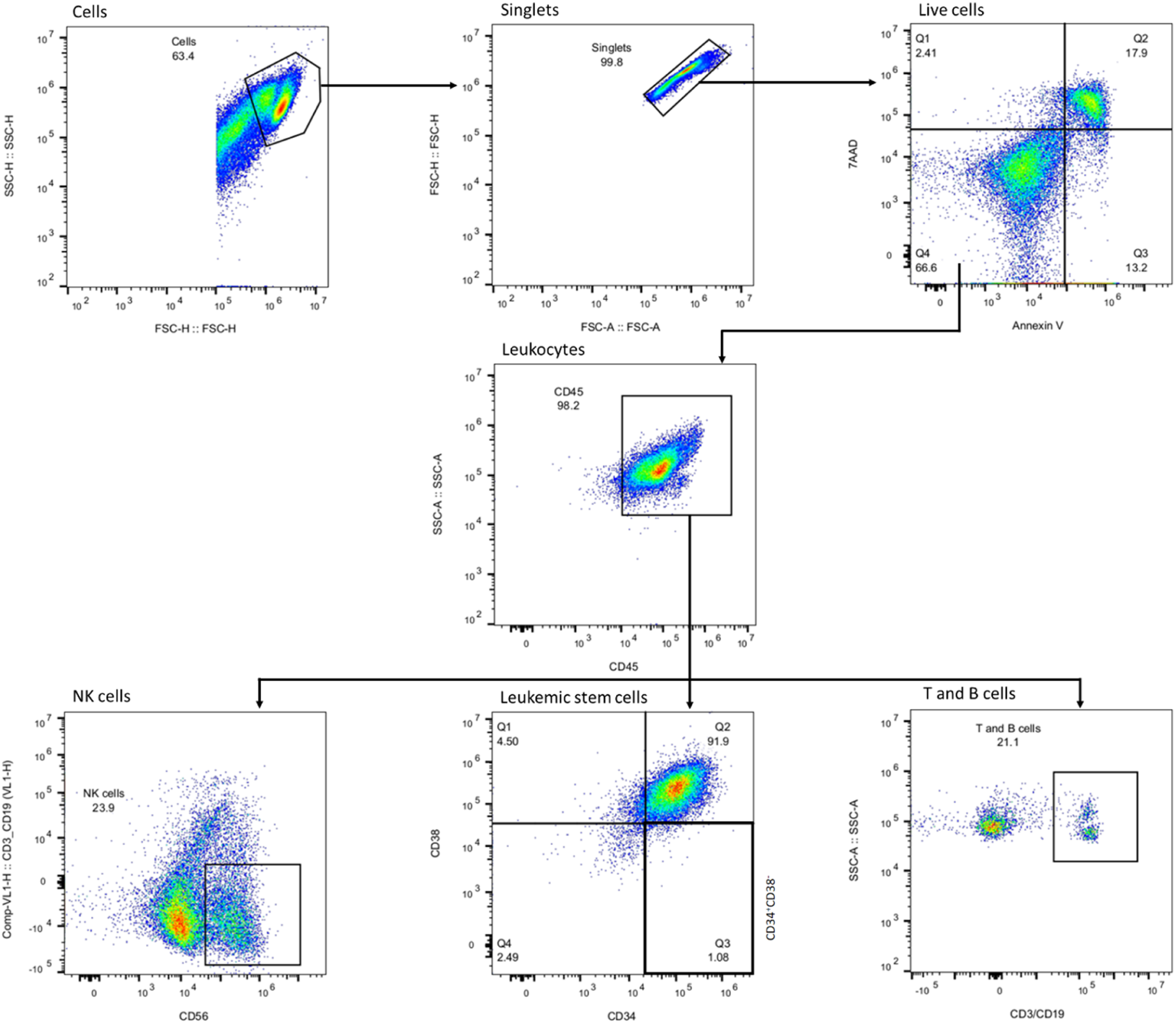
Gating strategy for the high-throughput flow cytometry (HTFC) assay. Live cells were gated as cells negative to 7-AAD and Annexin V markers. Leukemic stem cells were gated as CD34+ CD38-population. Lymphocytes were gated based on CD56 and CD3/CD19 to distinguish NK as well as T and B cell populations, respectively.

